# Chromatin conformation capture (Hi-C) sequencing of patient-derived xenografts: analysis guidelines

**DOI:** 10.1101/2020.10.17.343814

**Authors:** Mikhail G. Dozmorov, Katarzyna M. Tyc, Nathan C. Sheffield, David C. Boyd, Amy L. Olex, Jason Reed, J. Chuck Harrell

**Affiliations:** Department of Biostatistics, Virginia Commonwealth University, Richmond, VA, 23298, USA; Department of Pathology, Virginia Commonwealth University, Richmond, VA, 23284, USA; VCU Massey Cancer Center, Richmond, VA, 23298, USA; Center for Public Health Genomics, University of Virginia, Charlottesville, VA, 22908, USA; Integrative Life Sciences Doctoral Program, Virginia Commonwealth University, Richmond, VA, 23298, USA; C. Kenneth and Dianne Wright Center for Clinical and Translational Research, Virginia Commonwealth University, Richmond, VA, 23298, USA; Department of Physics, Virginia Commonwealth University, Richmond, VA, 23220, USA

**Author notes:** These authors contributed equally (Mikhail G. Dozmorov, Katarzyna M. Tyc).

**Keywords:** Hi-C, chromatin conformation capture, Xenografts, PDX, Xenome

## Abstract

Sequencing of patient-derived xenograft (PDX) mouse models allows investigation of the molecular mechanisms of human tumor samples engrafted in a mouse host. Thus, both human and mouse genetic material is sequenced. Several methods have been developed to remove mouse sequencing reads from RNA-seq or exome sequencing PDX data and improve the downstream signal. However, for more recent chromatin conformation capture technologies (Hi-C), the effect of mouse reads remains undefined.

We evaluated the effect of mouse read removal on the quality of Hi-C data using *in silico* created PDX Hi-C data with 10% and 30% mouse reads. Additionally, we generated two experimental PDX Hi-C datasets using different library preparation strategies. We evaluated three alignment strategies (Direct, Xenome, Combined) and three processing pipelines (Juicer, HiC-Pro, HiCExplorer) on the quality of Hi-C data.

Removal of mouse reads had little-to-no effect on data quality than the results obtained with Direct alignment strategy. Juicer pipeline extracted the most useful information from PDX Hi-C data. However, library preparation strategy had the largest effect on all quality metrics. Together, our study presents comprehensive guidelines on PDX Hi-C data processing.

## Introduction

Patient-derived tumor xenograft (PDX) mouse models are indispensable in preclinical and translational cancer research. Previous studies have demonstrated that human tumors engrafted in immunocompromised mouse models preserve each patient’s genetic heterogeneity (Bruna et al. 2016) and response to treatment (Izumchenko et al. 2017; DeRose et al. 2011). Consequently, the main application of PDX systems is to understand the molecular mechanisms of human cancers within controlled *in vivo* conditions. With the wide adoption of sequencing technologies, sequencing of PDX samples is now a standard (Turner et al. 2018; Alzubi et al. 2019; Girotti et al. 2016; Li et al. 2013).

High-throughput sequencing of PDX samples faces challenges not present when sequencing cell lines and homogeneous tissues. Engraftment of human cancer tissue fragments into mice leads to the rapid loss of human stroma and invasion of mouse stromal cells (Bruna et al. 2016; DeRose et al. 2011). Consequently, sequencing of PDX tumor samples produces reads derived from both human and mouse genomes, with mouse read contamination ranging from 4-7% up to 20% for RNA-seq and exome data (Rossello et al. 2013), and even 47% on average for whole-genome sequencing data (Lin et al. 2010). Metastases are even more variable, and we previously identified up to 99% mouse reads in PDX RNA-seq data from lung, liver, or brain metastases (Turner et al. 2018). Given the high similarity of human and mouse genomes, with orthologous gene products on average 85% identical (Makałowski et al. 1996), the presence of mouse reads introduces uncertainty in the alignment of PDX sequencing data.

Three strategies have been developed to address the removal of mouse reads from PDX sequencing data. The first strategy, referred hereafter as “Direct”, is the direct alignment of PDX sequencing data to the human genome. The second, filtering strategy, includes separation of human and mouse reads and using only human data for downstream analysis. The Xenome tool was among the first tools implementing filtering strategy. It classified reads into the human, mouse, both, neither, or ambiguous categories using a 25-mer matching algorithm (Conway et al. 2012). Despite being relatively old and lacking maintenance, Xenome remains widely used in bioinformatics pipelines (Woo et al. 2019). We refer to this strategy as “Xenome” throughout. The alignment of reads to human and mouse genomes and then filtering reads by best alignment match (Rossello et al. 2013) represents another viable approach to separate human and mouse reads. This approach has been implemented in Disambiguate (Ahdesmäki et al. 2016), bamcmp (Khandelwal et al. 2017), and XenoCP (Rusch et al. 2020) tools. The third strategy, referred hereafter as “Combined”, includes alignment to the *in silico* combined human-mouse reference genome to disambiguate human and mouse reads at the alignment step (Callari et al. 2018; Turner et al. 2018).

Each strategy for mouse read removal from PDX sequencing data has its own advantages and disadvantages. The filtering and combined strategies require extra efforts, more processing time, and in some cases doubling the requirements for computational resources. Several studies investigated the benefits of removal of contaminating mouse reads from PDX sequencing data. In DNA-seq PDX data, the removal of mouse reads reduced the false-positive rate of somatic mutation detection, especially when matching normal samples are not available (Woo et al. 2019; Rusch et al. 2020; Rossello et al. 2013; Ahdesmäki et al. 2016; Tso et al. 2014). In RNA-seq data, the removal of mouse reads improved gene expression quantification (Rusch et al. 2020), correlation with pure human gene expression (Rossello et al. 2013), and enrichment in relevant pathways (Khandelwal et al. 2017). Benchmarking of all three strategies using DNA sequencing convincingly demonstrated that filtering and combined strategies are necessary to minimize false discovery rates in detecting genomic variants, with exome sequencing data benefiting the most (Tso et al. 2014). The general consensus is that the removal of mouse reads from PDX sequencing data improves the extraction of human-specific signal from RNA-seq and DNA-seq PDX sequencing data (Conway et al. 2012; Callari et al. 2018; Rusch et al. 2020; Ahdesmäki et al. 2016; Khandelwal et al. 2017; Rossello et al. 2013; Woo et al. 2019).

Chromatin conformation capture technology and its high-throughput derivatives, such as Hi-C (Lieberman-Aiden et al. 2009), have recently emerged as tools to assess the three-dimensional (3D) structure of the genome. Changes in the 3D structure of the genome are an established hallmark of cancer (Rickman et al. 2012; Hnisz et al. 2016; Valton and Dekker 2016). However, the majority of the 3D cancer genomics studies have been performed *in vitro* using cell lines (Fritz et al. 2017; Johnston et al. 2019; Kantidze et al. 2020). Hi-C sequencing of PDX samples opens novel ways for understanding mechanisms of human cancers under controlled *in vivo* conditions. However, the effect of contaminating mouse reads on the quality of PDX Hi-C data, and the choice of the processing pipeline, remains undefined.

Hi-C sequencing data possesses unique qualities need to be considered when evaluating the effect of mouse reads in Hi-C PDX data. First, Hi-C paired-end reads are processed individually, as single-end data. Second, Hi-C data undergo extensive filtering to extract “valid pairs”, i.e., reads representative or two ligated DNA fragments with proper orientation and distance between them (Lajoie et al. 2015; Zheng et al. 2019). Furthermore, in contrast to typical sequencing experiments, processing of Hi-C data requires high-performance computational resources as one Hi-C experiment produces more than 20X number of reads of a typical RNA-seq experiment (Rao et al. 2014). It remains uncertain whether efforts for removing mouse reads from PDX Hi-C data are justified and meaningfully improve the quality of human Hi-C data.

To address the effect of mouse read removal in PDX sequencing data, we evaluated three strategies for preprocessing PDX Hi-C data: Direct, Xenome, and Combined. Using different library preparation strategies, we generated two deeply sequenced Hi-C datasets of a carboplatin-resistant UCD52 cell line (Turner et al. 2018; Alzubi et al. 2019). We further created three *in silico* PDX Hi-C datasets with either 10% or 30% of mouse read contamination, mirroring the percent of mouse reads observed in our experimental Hi-C data. In particular, we used Hi-C data from normal and cancer cells to investigate whether the biological properties, such as copy number variations inherent to cancer genomes, impact the quality of Hi-C data. Human Hi-C data without mouse reads contamination was used as a baseline. This design allowed us to comprehensively quantify the effect of contaminating mouse reads on the quality of Hi-C data and the downstream results.

Although several studies discuss how to process Hi-C data and what tools to use (Lajoie et al. 2015; Pal et al. 2019; Forcato et al. 2017), they have not evaluated the effect of mouse read contamination. We evaluated three leading pipelines, Juicer (Durand et al. 2016), HiC-Pro (Servant et al. 2015), and HiCExplorer (Ramirez et al. 2018) in terms of Hi-C data quality, extracted biological information, and computational runtime.

In total, we tested nine combinations of strategies–all pairwise combinations of three strategies for mouse read handling (Direct, Xenome, and Combined), and three processing tools (Juicer, HiC-Pro, and HiCExplorer)–to generate contact matrices from nine *in silico* and two experimental PDX Hi-C datasets. Furthermore, we assessed the effect of library preparation strategies on the quality of downstream results from Hi-C data. We found that removing mouse reads using Xenome or Combined strategies minimally affects the quality of Hi-C matrices and information extracted from them, while the Direct alignment yielded comparable-quality results without the additional computational overhead. The choice of processing pipeline showed detectable differences in data quality and the downstream results, with Juicer extracting the most information out of Hi-C data. Ultimately, the choice of library preparation strategy had the largest effect on data quality. From these studies, we recommend using the Direct alignment of PDX Hi-C data to the human genome. All three pipelines provided good quality results, with Juicer pipeline being our top choice. The choice of the library preparation strategy should be given a priority.

## Results

### A comprehensive workflow for assessing the impact of mouse read contamination in PDX Hi-C data

Sequencing of biological samples from patient-derived xenograft (PDX) mouse models face a challenge of mixed genomic context derived from host (mouse) and graft (human) cells. Naturally, the primary goal is to sequence human-specific genomic information; however, highly homologous mouse reads may hinder the identification of human genomic information. We investigated whether the presence of mouse reads in human Hi-C data negatively affects Hi-C data quality, and whether the removal of mouse reads improves the detection of topologically associating domains (TADs). Using the *in silico* and experimentally obtained PDX Hi-C data (Table 1, Supplementary Table S1), we assessed three alignment strategies for mouse read removal and three common pipelines to generate Hi-C matrices (Figure 1).

**Figure 1.**
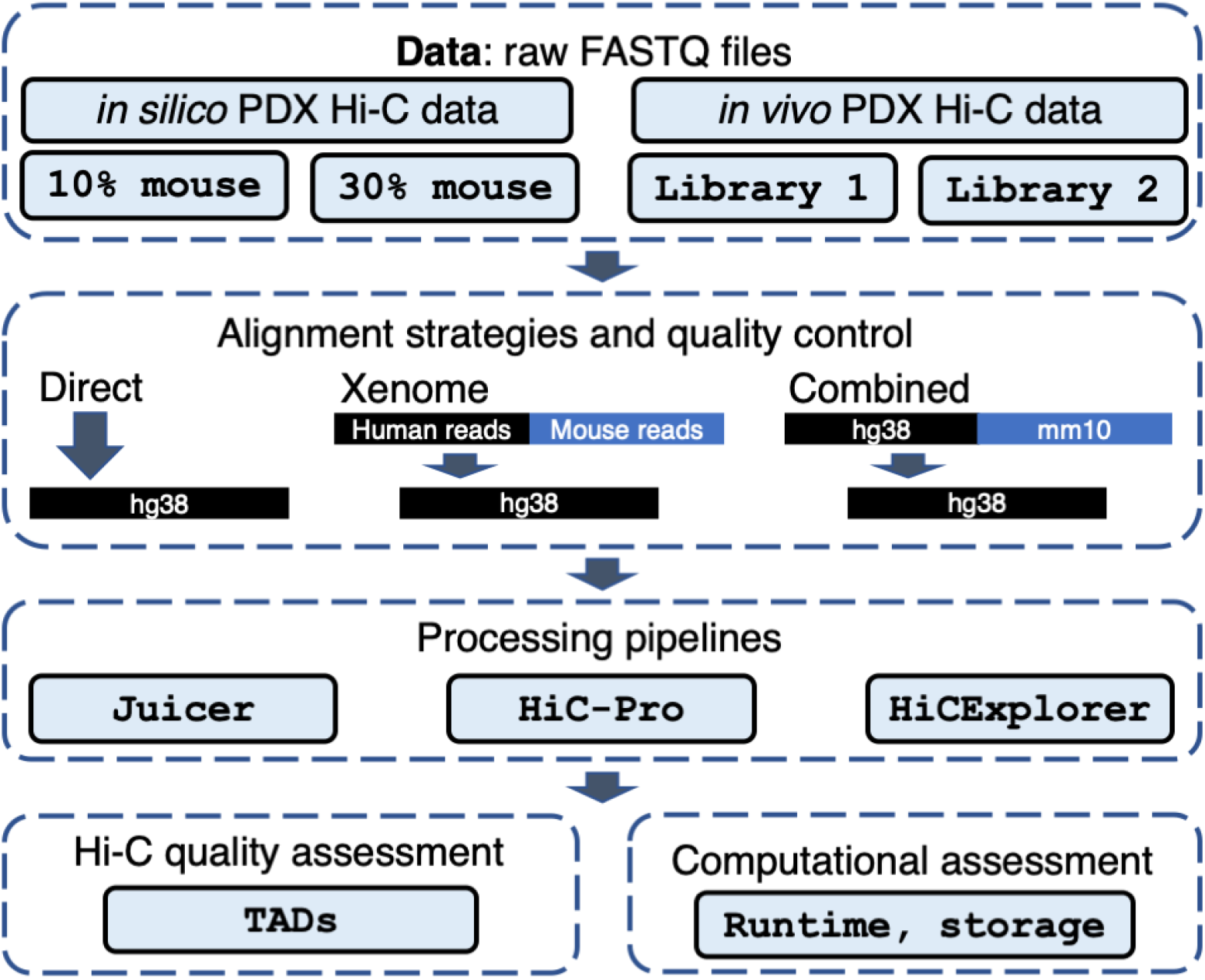
PDX Hi-C data analysis workflow. In silico (controlled mixture of human and 10/30% mouse Hi-C reads) and experimental PDX Hi-C data (two library preparation strategies) were processed using three read-alignment strategies (Direct: read alignment directly to the human genome, Xenome, and Combined: human reads retrieved with either Xenome or via pre-alignment step to the combined human-mouse genome, respectively). Three pipelines (Juicer, HiC-Pro, HiCExplorer) were used to obtain Hi-C matrices. Hi-C data quality and runtime metrics were assessed following each processing step.

**Table 1.** Summary of *in silico* and experimental PDX Hi-C data. The proportion of mouse reads within experimental PDX Hi-C data was estimated using Xenome/Combined alignment, respectively. The optimal resolution was estimated following Direct/Xenome/Combined alignment strategy, respectively.

| Hi-C data | Description | Total reads | Proportion of mouse reads | Optimal resolution (kb) |
| --- | --- | --- | --- | --- |
| **Baseline** |  |  |  |  |
| GM12878 | Human B-lymphoblastoids | 486,848,169 | 0% | 7.0 |
| HMEC | Human Mammary Epithelial | 456,577,383 | 0% | 7.9 |
| KBM7 | Human Myelogenous Leukemia | 431,368,621 | 0% | 8.3 |
| CH12-LX (rep 1) | Mouse lymphoma cell line | 45,594,869 | 100% | n/a |
| CH12-LX (rep 2) | Mouse lymphoma cell line | 175,930,719 | 100% | n/a |
| **in silico** |  |  |  |  |
| **PDX** |  |  |  |  |
| GM12878 (10%) | GM12878 + CH12-LX (rep 1) | 532,443,038 | 8.56% | 7.0/7.1/7.0 |
| GM12878 (30%) | GM12878 + CH12-LX (rep 2) | 662,778,888 | 26.54% | 7.0/7.1/7.0 |
| HMEC (10%) | HMEC + CH12-LX (rep 1) | 502,172,252 | 9.08% | 7.9/7.9/7.9 |
| HMEC (30%) | HMEC + CH12-LX (rep 2) | 632,508,102 | 27.81% | 7.9/7.9/7.9 |
| KBM7 (10%) | KBM7 + CH12-LX (rep 1) | 476,963,490 | 9.56% | 8.3/8.3/8.3 |
| KBM7 (30%) | KBM7 + CH12-LX (rep 2) | 607,299,340 | 28.97% | 8.3/8.3/8.3 |
| **Experimental** |  |  |  |  |
| **PDX** |  |  |  |  |
| UCD52 Library 1 | Basal-like BRCA cell line | 873,892,191 | 12.16%/12.38% | 11.5/11.9/11.7 |
| UCD52 Library 2 | Basal-like BRCA cell line | 708,069,622 | 25.78%/29.14% | 8.9/9.1/9.0 |

The *in silico* PDX Hi-C data were created by concatenating previously published mouse and human Hi-C data (Rao et al. 2014) (see Methods). Human Hi-C data from GM12878 B-lymphoblastoid cells (nearly normal karyotype) and KBM7 myelogenous leukemia (near-haploid karyotype) were selected to assess the effect of mouse read contamination in normal and cancer Hi-C data, respectively. HMEC human mammary epithelial cells were selected to parallel breast cancer origin of our experimental PDX Hi-C data. Mouse Hi-C data from B-lymphoblast CH12-LX cells were used to create the *in silico* PDX Hi-C data with ~ 10% and ~ 30% level of mouse read contamination. Human Hi-C data for the corresponding cell lines without mouse reads were used as a baseline.

The main limitation of *in-silico* PDX Hi-C data is that human and mouse reads originate from different libraries. Although *in silico* PDX Hi-C data may be sufficient to test the performance of aligners on a mixture of highly homologous human and mouse reads, it is unknown whether this mixture can recapitulate the complexity of experimental PDX Hi-C data, where, theoretically, crosslinking and ligation of human and mouse DNA can occur. To investigate whether the removal of mouse reads from experimental PDX Hi-C data improves the quality of Hi-C matrices, we generated replicates of Hi-C data from a triple-negative breast cancer PDX (UCD52 cells), obtained with two different library preparation strategies (Library 1 and Library 2, see Methods). As expected, human-specific replicates of experimental PDX Hi-C data prepared with the same library preparation strategy showed high correlation, in contrast to those prepared awith different strategy (average Pearson Correlation Coefficient PCC = 0.9963 and 0.9547, respectively). Mouse matrices were uniformly correlated irrespective of the library preparation strategy (average PCC = 0.9870, Supplementary Figure S1). Therefore, replicates of Hi-C data were merged for downstream processing. In total, we processed 11 PDX Hi-C datasets (Table 1).

We applied three alignment strategies to remove mouse reads contamination: the Direct alignment of PDX Hi-C reads to the human reference genome (“Direct”), the alignment of data cleaned of mouse reads data using the Xenome tool (Conway et al. 2012) (“Xenome”), or using pre-alignment to a combined human and mouse genomes (“Combined”, see Methods, Figure 1). Three tools for processing of Hi-C data were applied: Juicer (Durand et al. 2016), HiC-Pro (Servant et al. 2015), and HiCExplorer (Ramirez et al. 2018) (Figure 1). The use of different methods for mouse read removal and processing pipelines allowed for finding the optimal strategy for analyzing Hi-C data derived from PDX mouse models.

### Experimental PDX Hi-C data have higher proportion of ambiguously mapped reads

Xenome accurately estimated the 10%/30% proportion of mouse reads in our *in silico* PDX Hi-C data (Figure 2, Supplementary Table S2). We observed a similar proportion of mouse reads in our experimental PDX data (approximately 12% and 30%, Table 1). Less than 1% of reads were mapped to both or neither human nor mouse genomes, and these results were consistent in the *in silico* and experimental PDX Hi-C data. Compared with *in silico* PDX data, the number of “ambiguous” reads in the experimental data was slightly higher (4-5% vs. 1%, Supplementary Table S2). Overall, our results indicate that *in silico* PDX Hi-C data reflect the level of mouse reads contamination observed in experimental settings. However, the higher level of ambiguously mapped reads suggests unique biological properties in experimental PDX Hi-C data and justifies the need for its analysis.

**Figure 2.**
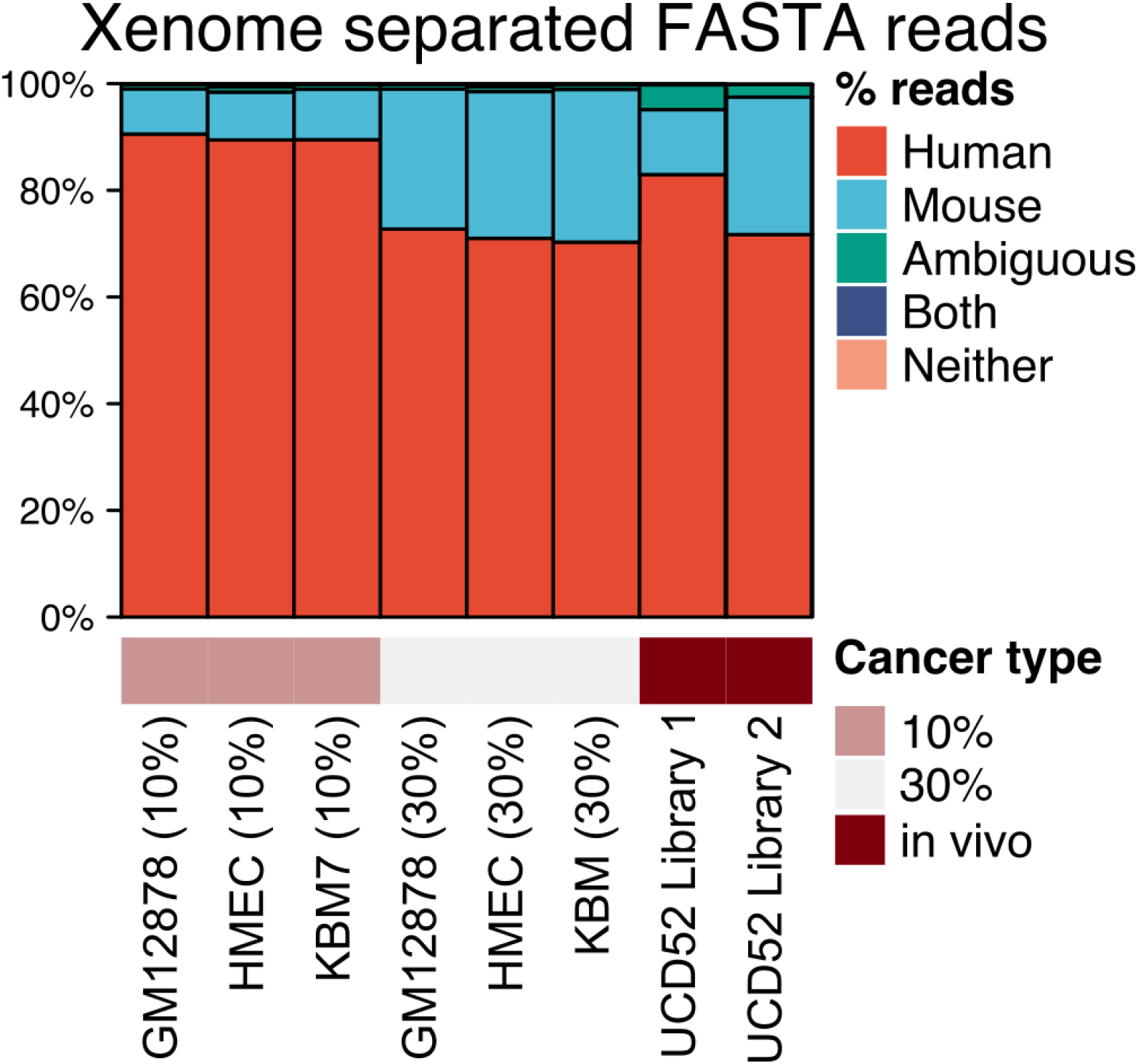
Proportions of human and mouse reads in experimental and *in silico* PDX Hi-C data. Details of Xenome read separation statistics are shown in Supplementary Table S2.

### Removal of mouse reads has negligible impact on Hi-C contacts retrieval rate and their quality

Following data processing using all combinations of alignment strategies and downstream tools, we extracted four Hi-C quality metrics from the log files produced by each pipeline (all QC metrics are shown in Supplementary Table S3). **Alignment rate** is the proportion of reads aligned to the human genome. **Valid interaction pairs** is the proportion of reads marked as Hi-C contacts by each tool considering the valid restriction site within a reasonable distance. Higher values of those metrics indicate better data quality. **Cis/trans ratio** is the ratio of intra- vs. inter-chromosomal interacting reads. A higher cis/trans ratio indicates an enrichment for within-chromosomal reads, expected in the Hi-C experiments. **Long/short ratio** is the ratio of cis interactions more than 20kb away vs. those less than 20kb away. The expectation is to capture more long-distance chromatin interactions, i.e., a long/short ratio with a value higher than 1, while the long/short ratio less than 1 indicates long interactions are lost. These Hi-C quality metrics allow for the comprehensive definition of optimal alignment strategy and the effect of mouse read removal.

The removal of mouse reads had minimal-to-no effect on the alignment quality metrics of *in silico* and experimental PDX Hi-C data (Figure 3, Supplementary Figure S2). Expectedly, the alignment rate and the proportion of valid interaction pairs in *in silico* PDX Hi-C data were diminished proportionally to the percent of mouse read contamination (10% or 30%), as compared with those in pure human Hi-C data for the corresponding cell lines (Figure 3A, B). The removal of mouse reads from *in silico* PDX Hi-C data did not markedly affect the cis/trans ratio and long/short ratio (Figure 3C, D). These results were consistent across cell lines (Supplementary Figure S2). These results suggest that the Direct alignment strategy of *in silico* PDX Hi-C data performs similarly to using data with explicitly removed mouse reads.

**Figure 3.**
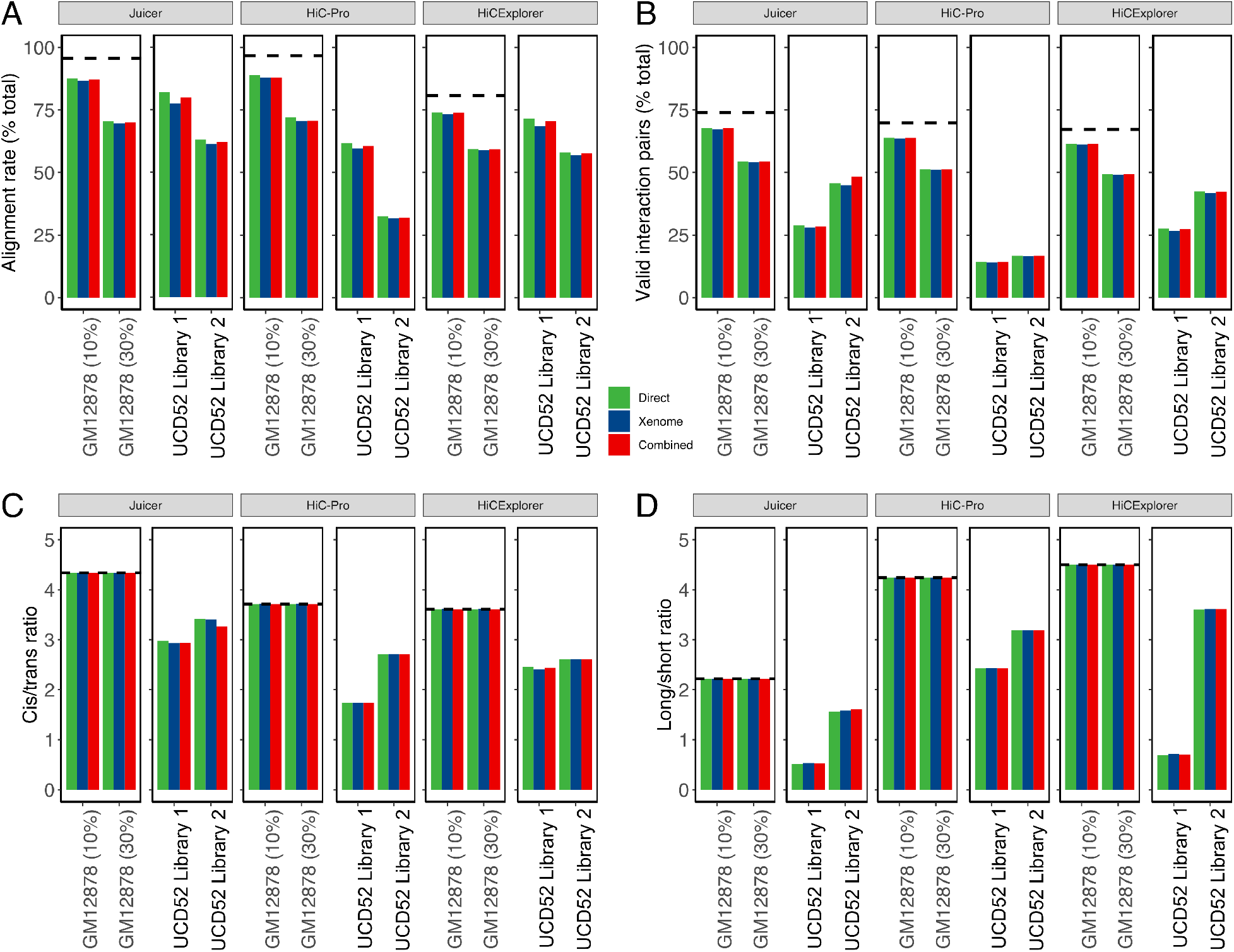
Quality metrics for selecting the optimal PDX Hi-C data processing pipeline. All metrics are stratified by the pipeline (Juicer, HiC-Pro, and HiCExplorer) and color-coded by the alignment strategy (Green: Direct, Blue: Xenome, Red: Combined). (A) Alignment rate representing the proportion of all aligned reads. (B) The proportion of valid interaction pairs as determined by each pipeline. (C) The ratio of Cis interacting pairs (i.e., occurring on the same chromosome) vs. trans interacting pairs (i.e., between chromosome interactions). (D) The ratio of long-vs. short-interacting Hi-C contacts. Dashed lines correspond to the baseline alignment quality metrics for Hi-C data without mouse reads.

Similar to the results obtained with *in silico* PDX Hi-C data, the removal of mouse reads from experimental PDX Hi-C data did not markedly affect quality metrics (Figure 3), although more variability was observed (~2-4%). Interestingly, although the alignment rate of data prepared with Library 2 strategy was lower than that of Library 1-prepared data (Figure 3A), the proportion of valid interaction pairs, cis/trans ratio, and, in particular, long/short ratio were higher (Figure 3B-D). These results suggest that the Library 2-prepared data contain more information about intra-chromosomal long- and short-distance chromatin interactions. In summary, these results indicate that the removal of mouse reads does not substantially improve or change the alignment quality of PDX Hi-C data, but the library preparation strategy has a significant effect.

### Juicer pipeline recovers more useful information from PDX Hi-C data

Although removing mouse reads using either strategy did not substantially affect the alignment quality of PDX Hi-C data, we noted pipeline-specific differences (Figure 3, Supplementary Figure S3), referred by their names for brevity. Specifically, Juicer produced a similar alignment rate as HiC-Pro in *in silico* PDX Hi-C data. However, it recovered nearly 15% more alignable reads in experimental PDX Hi-C data. Compared with Juicer and HiC-Pro, HiCExplorer yielded ~20% lower alignment rate for *in silico* PDX Hi-C data. Yet, HiCExplorer performed well and only slightly inferior to Juicer for the alignment of experimental PDX Hi-C data (Supplementary Figure S3A). Similarly, Juicer recovered up to a 10% higher proportion of valid interaction pairs in *in silico* PDX data than HiC-Pro and HiCExplorer (Supplementary Figure S3B). However, in experimental PDX Hi-C data, Juicer recovered nearly twice as many valid interaction pairs as the HiC-Pro, and outperformed HiCExplorer by ~2% margin (Supplementary Figure S3B). These results indicate that Juicer can recover more alignable reads and recover a higher proportion of valid interaction pairs. These improvements were particularly pronounced when processing experimental PDX Hi-C data.

A typical Hi-C experiment is expected to detect the majority of interactions within chromosomes (cis interactions) as compared with between chromosome (trans) interactions. This should be reflected by a high cis/trans ratio. Juicer produced Hi-C data with a higher cis/trans ratio than HiC-Pro and HiCExplorer pipelines. These results were consistent between *in silico* and experimental PDX Hi-C data (Figure 3, Supplementary Figure S3C). Importantly, Juicer yielded slightly lower long/short ratios compared to other two pipelines (Figure 3D), which reflects the fact that Juicer captured more cis interacting interaction (Figure 3C). These results were consistent in *in silico* and experimental PDX Hi-C data (Supplementary Figure S3D). Notably, all quality metrics were superior in Hi-C data obtained using the Library 2 preparation strategy. These results confirm the better performance of Juicer for extracting the maximum amount of useful information from PDX Hi-C data.

### Juicer pipeline better recovers short- and long-distance interactions

The frequency of chromatin interactions follows a power-law decay with the increased distance between interacting regions (Lajoie et al. 2015). Subsequently, the power-law exponent is an indicator of the rate of the decay, with smaller values corresponding to slower decay due to the presence of long-range interactions. Using the Direct alignment strategy, we investigated the effect of mouse reads and the pipeline choice on the distance-dependent decay of interaction frequencies. The presence of mouse reads did not affect the rate of the decay, and these results were consistent across PDXs (Figure 4A, Supplementary Figure S4, Supplementary Table S4). However, we noted that the pipeline choice affected the power-law decay, and this effect was most pronounced in experimental PDX Hi-C data (Figure 4A, Supplementary Table S4). Juicer-processed data showed a consistently smaller exponent of the power-law decay of chromatin interaction frequencies (Figure 4BC, Supplementary Table S4). Experimental PDX Hi-C data obtained with the Library 2 preparation strategy showed, on average, smaller and more stable power-law exponent (1.83 ± 0.01) than that of Library 1-prepared data (1.99 ± 0.22, Supplementary Table S4). These results suggest that Juicer better preserves short- and long-distance interactions leading to a slower decrease in the average interaction frequency. In summary, our results indicate that Juicer performs best for extracting maximum information from PDX Hi-C data, and the choice of library preparation strategy is critical for optimal data quality.

**Figure 4.**
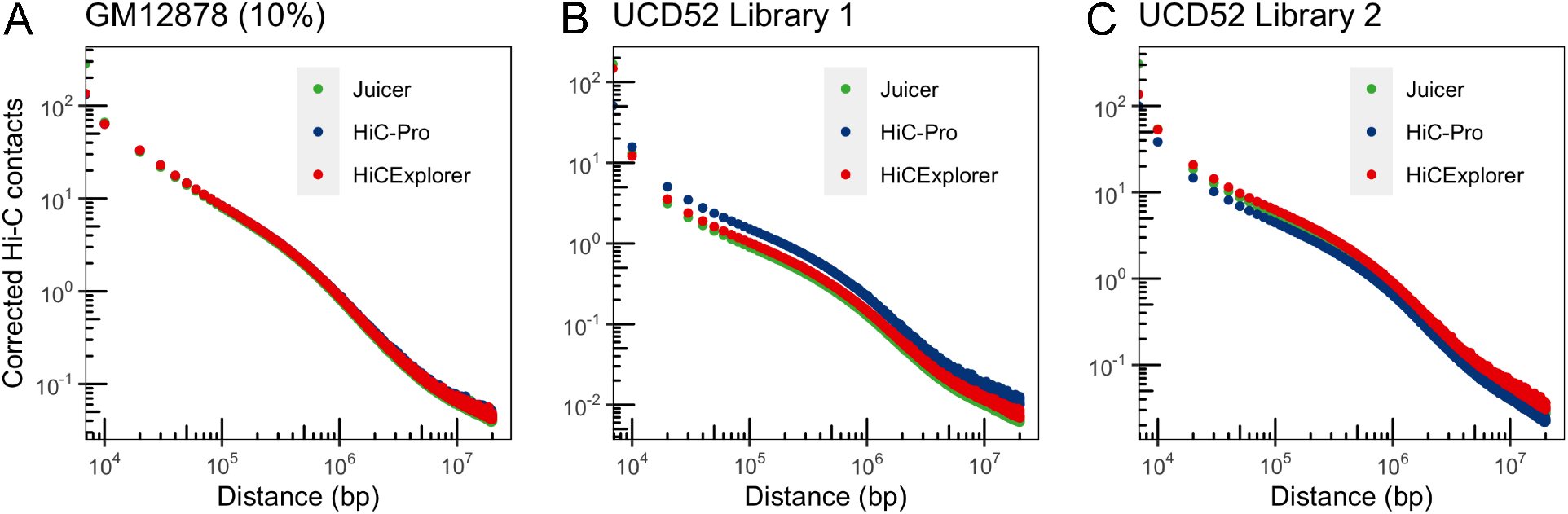
Slower rate of the distance-dependent decay of chromatin interactions in Juicer-processed data. The rate of distance-dependent decay in the presence of mouse reads is illustrated with *in silico* PDX Hi-C data using GM12878 cell line with 10% mouse reads (A) and compared to two experimental PDX Hi-C data (B, C). Pipelines used to process the data are indicated in the color legend. Results for the Direct alignment strategy are shown.

### The presence of mouse reads has a negligible effect on the detection of Topologically Associating Domains

The most typical use of Hi-C data is to detect common 3D structures, such as topologically associating domains (TADs). Using the Direct alignment strategy, we evaluated the number and size of TADs detected from data processed by the three pipelines. Compared to the baseline (pure human Hi-C data), the number of cell-type-specific TADs was nearly identical at the 10% or 30% level of *in silico* mouse read contamination (Figure 5A, Supplementary Table S5). These results were consistent irrespectively of the pipeline. These results indicate that the presence of mouse reads in PDX Hi-C data has minimal effect on the number of called TADs.

**Figure 5.**
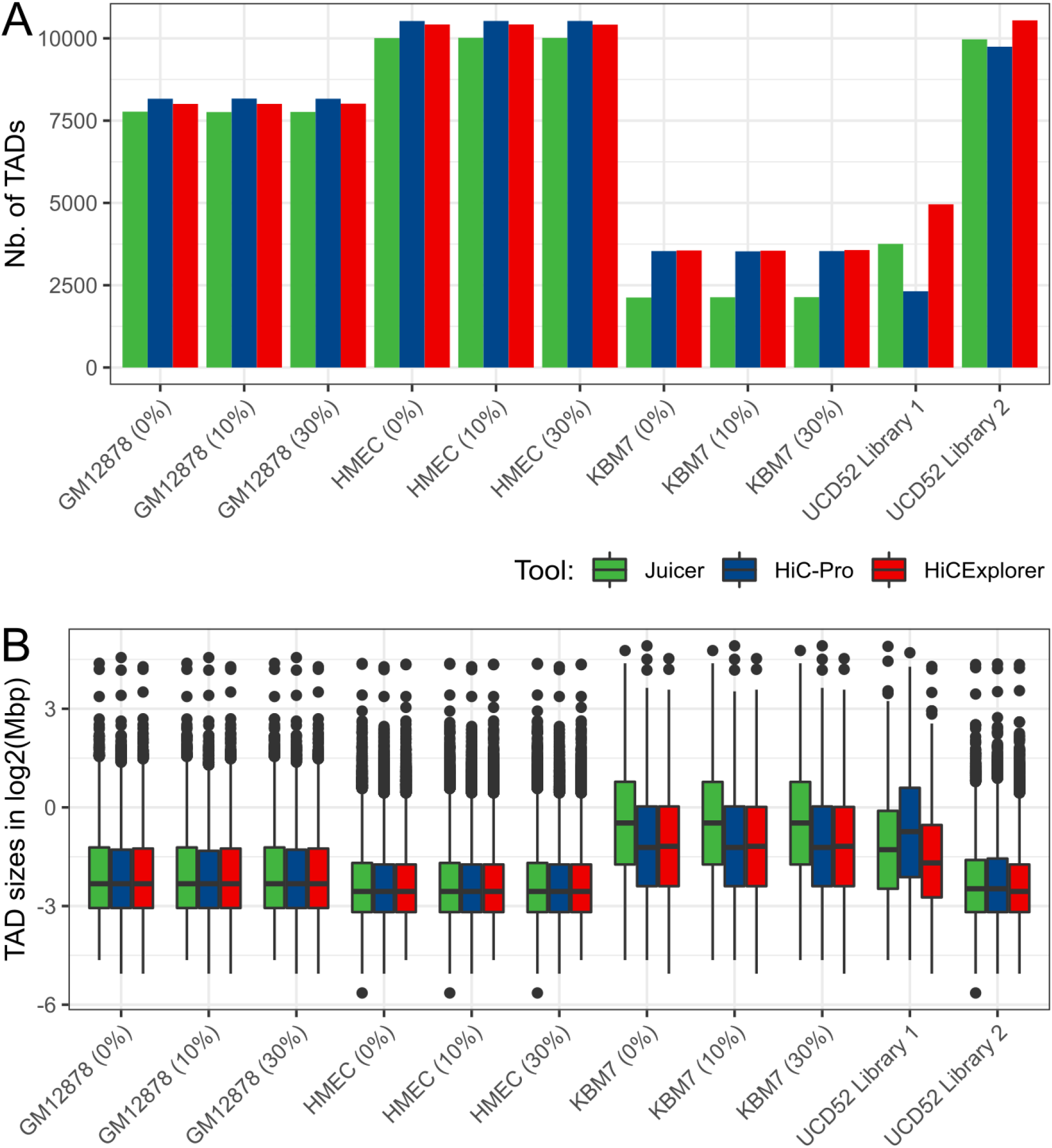
The library preparation strategy has the largest effect on TAD detection. The number (A) and the width of TADs (B) are similar at different levels of mouse reads and across processing pipelines in all but Library 1-prepared experimental PDX Hi-C data. Results for the Direct alignment strategy are shown.

### Juicer-processed data contains fewer TADs

While mouse read contamination did not markedly affect the number of TADs, the choice of processing pipeline had a variable effect on the number of TADs. Expectedly, the total number of TADs differed between cell lines used for *in silico* PDX Hi-C data construction and library preparation strategies for experimental PDX Hi-C data (Figure 5A). We observed a smaller number of TADs detected in Juicer-processed *in silico* data, paralleled by the larger TAD size (Figure 5B). However, in experimental PDX Hi-C data, HiC-Pro-processed data had the smallest number of TADs, followed by Juicer-processed data. These differences were less pronounced in Library 2-prepared data, where more than twice as many TADs were detected as compared with the Library 1-prepared data (Figure 5A). In summary, TADs detected from Juicer-processed data tend to be wider, an effect less pronounced compared to the library preparation strategy effect.

### The choice of library preparation strategy has the largest effect on TAD detection

We observed nearly twice as many TADs detected in Library 2-prepared data than those detected in Library 1-prepared data (Figure 5A). Consequently, the size of TADs detected in Library 2-prepared data was smaller (Figure 5B). These results parallel our observation that Library 2-prepared data has better quality metrics (Figure 3); however, the recovery of more than twice as many TADs was unexpected. Notably, the pipeline-specific differences in TAD number and their size detected in Library 2-prepared data were negligible, similar to what we observed for the *in silico* PDX Hi-C data. This is in contrast to Library 1-prepared data, where the differences in pipelines were more pronounced (Figure 5). These results suggest that, for the optimal library preparation strategy, the differences in pipelines are negligible, emphasizing the importance of library preparation strategy.

### Technical and runtime considerations

We compared the runtime and storage requirements for each alignment strategy and pipeline. Removal of mouse reads with either Xenome or Combined strategy resulted in smaller datasets and, consequently, faster processing time (Figure 6A). However, when considering the additional time needed to remove mouse reads (longest for the Combined strategy), processing of the original data was the fastest. Together with previous observations of the minimal effect of mouse read removal on Hi-C data quality, these results indicate that extra computational time used to remove mouse reads may not be beneficial for the quality of downstream results.

**Figure 6.**
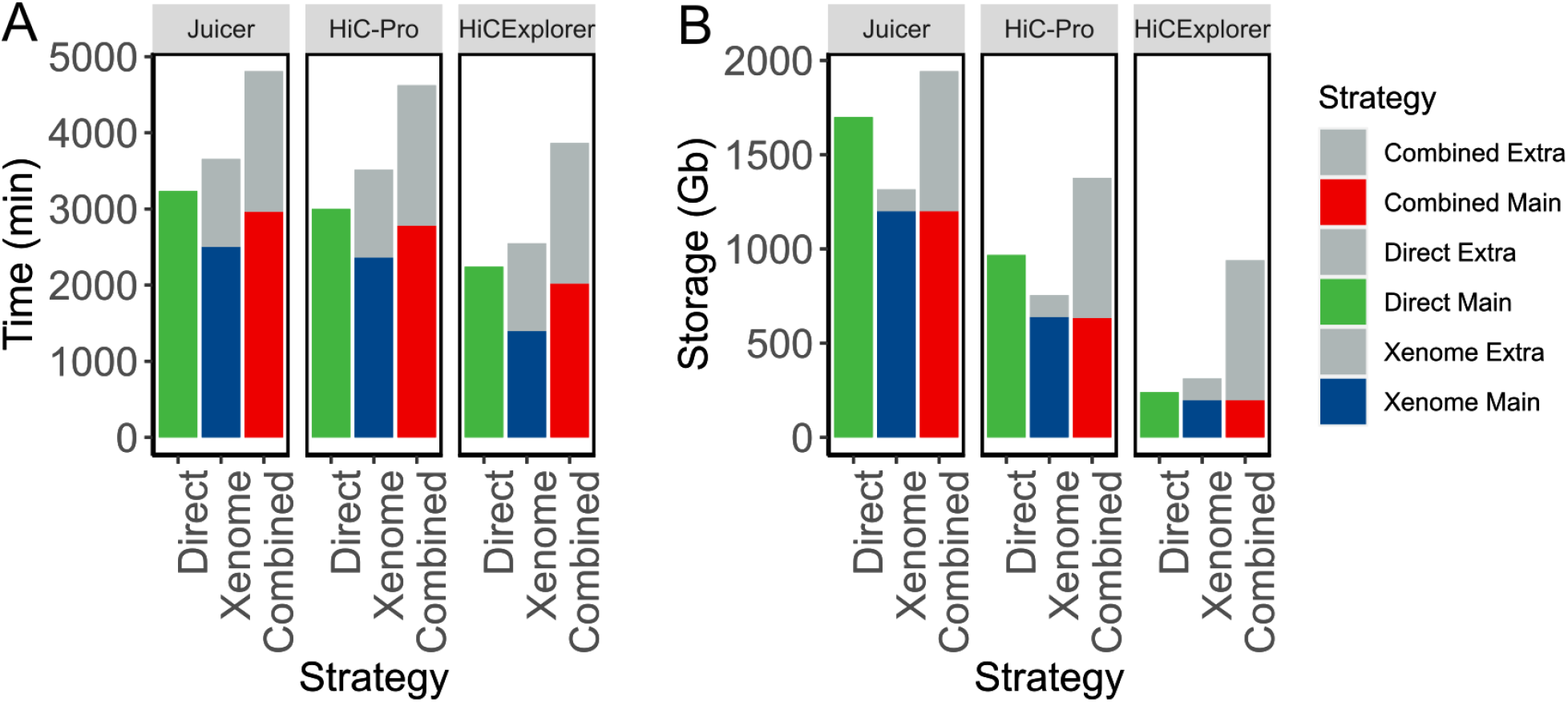
Removal of mouse reads carries a significant computational overhead. An example of runtime (A) and storage (B) resources required for processing PDX Hi-C data to obtain matrices. Only within-pipeline runtime comparisons are valid, as pipelines used different computational resources (see Methods). Results for processing Library 2-prepared PDX Hi-C data are presented. Extra: accounting for time and storage space required to filter mouse reads. Main: time and storage determined for processing human reads.

The removal of mouse reads requires extra storage space, with the Combined strategy requiring the largest amount of additional storage (Figure 6B). Interestingly, the Juicer pipeline required the largest storage space; however, it can be minimized by compressing text files produced by it. Expectedly, processing of smaller PDX Hi-C data without mouse reads required less space. Together with additional time requirements, extra space for removing mouse reads creates a significant computational overhead with negligible benefits as compared with the Direct alignment strategy.

The choice of tools for mouse read removal is an important technical consideration requiring significant human time. Xenome, a part of the Gossamer bioinformatics suite, has not been updated since January 5, 2017 (as of October 15, 2020). It requires dependencies that can only be installed using administrative privileges, which are rarely available for bioinformaticians working in a high-performance computing environment. Furthermore, Xenome requires the creation of its own genome index, which also contributes to the storage and processing time, and was not included in Figure 6. The Combined strategy can be implemented ad hoc, and the combined genomes and indexes can be downloaded using Refgenie (Stolarczyk et al. 2020) (see Methods). However, the extra hard drive space and time required for mouse read removal create an unnecessary human and computational burden and can contribute to delays in a project. Take together, we recommend using the Direct alignment strategy and Juicer pipeline for the most optimal computational processing of experimental PDX Hi-C data.

## Discussion

We have assessed the impact of mouse read contamination on the performance of three leading pipelines for Hi-C data processing. Using quality control metrics at the alignment stage, we showed that, unlike whole-exome and RNA-seq data from PDX models, Hi-C PDX data are unaffected by mouse read contamination. This is not unexpected as Hi-C data processing pipelines include a series of filters to select valid pairs (Lajoie et al. 2015). It is highly unlikely for experimental PDX Hi-C data to contain human-mouse chimeric reads, and even if such a read pair occurs, the probability that it would be recognized as a valid Hi-C contact (i.e., mapped in the proper orientation, within a certain distance from the nearest restriction site, etc.) is negligible. Results from our study confirm this reasoning and recommend the Direct alignment of PDX Hi-C data to the graft (human) genome.

Our results indicate that the Juicer pipeline consistently recovers more alignable reads, valid interaction pairs, and achieves better cis/trans and long/short interaction ratios. The better performance of Juicer over the HiC-Pro pipeline can be attributed to the use of the `bwa mem` algorithm that can efficiently handle split-read alignment. In contrast, HiC-Pro uses `bowtie2` aligner with the default “–end-to-end” mapping settings. However, despite HiCExplorer uses `bwa mem` aligner, its performance was marginally inferior to that of Juicer. Our results are in line with recommendations to use `bwa mem`-based pipelines when processing experimental PDX Hi-C data, with Juicer being the better pipeline for extracting most useful data.

The library preparation strategy appeared to play a major role in downstream data quality. While differences in quality metrics between *in silico* PDX Hi-C datasets can be attributed to the differences in sequencing depth (Supplementary Table S1), differences in our experimental PDX Hi-C data can be directly attributed to the library preparation strategies. Although our experimental PDX Hi-C data had nearly twice as many reads as the *in silico* PDX Hi-C data (Table 1), their quality metrics were inferior compared to *in silico* constructed Hi-C data (Figure 3). This was most pronounced for Library 1-prepared data, which we speculate is due to the presence of nearly 40% read duplicates, as compared to 12-15% duplicates in other datasets (Supplementary Table S1). However, the higher proportion of dangling ends, self circles, dumped reads, singletons etc. may have contributed to the inferior quality of Library 1-prepared data (Supplementary Table S3). Similar to the ENCODE guidelines (ENCODE project 2018), our observations suggest the importance of controlling duplicates in Hi-C data and.

Despite the lower number of sequencing reads and alignment rate, data obtained with Library 2 preparation strategy recovered more cis interacting Hi-C contacts spanning longer distances (cis/trans ratio and long/short ratio metrics in Figure 3C and Figure 3D, respectively), and slower distance-dependent decay (Figure 4, Supplementary Table S4). Furthermore, the number and size of TADs detected from the Library 2-prepared data was similar to that of detected in *in silico* PDX Hi-C data (Figure 5). This can be attributed to multiple-enzyme genome digestion that cut the human genome in more than 16M sites. In contrast, the single-enzyme Library 1 preparation strategy digests the genome in about 7.2M sites. Given Hi-C data quality significantly affects downstream results, we suggest careful inspection of the shallow sequencing data prior to the deep-sequencing experiment, giving particular weight to the metrics presented in Figure 3. The choice of restriction enzymes should be given the primary consideration in designing PDX Hi-C experiments.

According to the ENCODE guidelines (ENCODE project 2018), we expected to recover about 58% of sequenced reads as valid Hi-C interactions. While our *in silico* PDX Hi-C data (Rao et al. 2014) almost always achieved this threshold, our experimental PDXs did not meet these criteria (~ 28 and ~ 45 for Library 1 and Library 2 preparation strategies, respectively, Supplementary Table S3). Of note, other studies report a much lower rate of valid Hi-C interactions. For instance, the average number of valid interactions across 93 Hi-C datasets was 17.72 ± 13.04 (Yang et al. 2018). The overall lower percentage of valid interactions in our experimental Hi-C data can be partially explained by the fact that the genome of carboplatin-resistant UCD52 cells may be affected by genome rearrangements. The presence of duplications, deletions, and inversions is known to affect the genome’s 3D organization (Chakraborty and Ay 2017) and may have negatively affected the performance of our experimental PDX Hi-C data. Our results suggest the need to consider the effect of large-scale genome variation in the processing of PDX Hi-C data, in addition to the standard Hi-C data quality metrics.

## Methods

### Generation of experimental PDX Hi-C data

UCD52 tumors were implanted in mice and once palpable treated with a single dose of 40mg/kg carboplatin, as previously described (Turner et al. 2018; Alzubi et al. 2019). Once the tumors began growing again, they were treated with another dose of carboplatin. This was repeated until the tumor was no longer responsive to carboplatin. Xenograft tissue samples were processed by Phase Genomics (Seattle, WA) and Arima Genomics (San Diego, CA). Data generated using Phase Genomics/Arima Genomics library preparation strategy are referred to as “Library 1”/“Library 2”, respectively. The following protocols detail each strategy, as provided by the respective service providers.

### Phase Genomics (Library 1) preparation strategy

Approximately 200 mg of xenograft tissue was finely chopped and then crosslinked for 20 min at room temperature (RT) with end-over-end mixing in 1 ml of Proximo Crosslinking solution. The crosslinking reaction was terminated with a quenching solution for 20 min at RT with end-over-end mixing. Quenched tissue was rinsed once with 1X Chromatin Rinse Buffer (CRB), resuspended in Proximo Animal Lysis Buffer 1, and then transferred to Dounce Homogenizer (Kontes) and homogenized with 12 strokes using the ‘A’ homogenizer. Disrupted tissue in lysis buffer was incubated 20 min at RT. Large debris was removed following a 1 min 500xg spin. Lysate was recovered and transferred to a clean tube and pelleted by spinning at 17,000xg for 5 min. The supernatant was removed and pellet washed once with 1X CRB. After removing 1X CRB wash, the pellet was resuspended in 100 *μl* Proximo Lysis Buffer 2 and incubated at 65°C for 10 min Chromatin was irreversibly bound to SPRI beads by adding 100 *μl* SPRI beads to lysate, incubating for 10 min at RT. Beads were then washed once with 1X CRB. Beads were resuspended in 150 *μl* of Proximo fragmentation buffer and 5 *μl* of Proximo fragmentation enzyme (PN LS0027; 5,000 U/ml Sau3AI cutting at ‘GATC’) was added and incubated for 1 hour at 37°C. The sample was cooled to 12°C, and 2.5 μl of Phase Genomics Finishing Enzyme was added (PN LS0030). Sample was incubated 30 minutes at 12°C, adding 6 μl of Stop Solution (PN LS0004) at the completion of the incubation. The beads were then washed with 1X CRB and resuspended in 100 μl of Proximo Ligation Buffer supplemented with 5 μl of Proximity ligation enzyme. The proximity ligation reaction was incubated at RT for 4 hours with occasional gentle mixing. After the ligation step, 5 μl of Reverse Crosslinks enzyme (PN BR0012) was added and the reaction incubated at 65°C for 1 hour. After reversing crosslinks, the free DNA was recovered by adding 100 μl of SPRI buffer to the reaction. Beads were washed twice with 80% ethanol, air dried, and proximity ligation products were eluted (Elution Buffer, PN BR0014). DNA fragments containing proximity ligation junctions were enriched with streptavidin beads (PN LS0020). After washing streptavidin beads twice with PG Wash Buffer 2 (PN BR0004), once with PG Wash Buffer 1 (PN BR0016), and once with molecular biology grade water, library was constructed using Proximo library reagents (PNs LS0034, LS0035, and BR0017) amplified with high-fidelity polymerase (PN BR0018), and size selected using SPRI enriching for fragments between 300 and 700 bp. Pooled libraries were sequenced on an Illumina NovaSeq 6000 instrument using an S4 flow cell. Libraries were de-multiplexed using unique dual indexes following standard Illumina methods.

### Arima Genomics (Library 2) preparation strategy

Hi-C experiments were performed by Arima Genomics (https://arimagenomics.com/) according to the Arima-HiC protocols described in the Arima-HiC kit (P/N : A510008). After the Arima-HiC protocol, Illumina-compatible sequencing libraries were prepared by first shearing purified Arima-HiC proximally-ligated DNA and then size-selecting DNA fragments from ~200-600bp using SPRI beads. The size-selected fragments were then enriched for biotin and converted into Illumina-compatible sequencing libraries using the KAPA Hyper Prep kit (P/N: KK8504). After adapter ligation, DNA was PCR amplified and purified using SPRI beads. The purified DNA underwent standard QC (qPCR and Bioanalyzer) and was sequenced on the HiSeq X following the manufacturer’s protocols.

### Construction of *in silico* PDX Hi-C data

Publicly available Hi-C data from Rao et al. 2014 study (Rao et al. 2014) (GSE63525) were used to construct *in silico* PDX Hi-C data. Three human and one mouse cell line Hi-C data were selected (Table S1). To construct *in silico* PDX data containing a mixture of human and mouse reads, FASTA files from human and mouse cell lines were concatenated to form Hi-C datasets containing approximately 10% and 30% mouse reads (Table 1). If read length differed between human and mouse datasets, reads were trimmed from 3’ end to 96bp (smallest read length) using cutadapt (v2.7, (Martin 2011)) before concatenation.

### Removal of mouse reads from PDX Hi-C data

Three mouse read removal strategies were evaluated: Direct, Xenome, and Combined (Figure 1). In the Direct alignment strategy, all reads were mapped to the human reference genome version GRCh38/hg38 using only autosomal and sex chromosomes. In the Xenome approach, PDX Hi-C reads were processed with the `Xenome` tool (Conway et al. 2012) from the gossamer GitHub repository (https://github.com/data61/gossamer), and human only FASTA reads were kept. In the Combined strategy, the combined human-mouse genome was created by concatenating autosomal and sex chromosomes from hg38 and mm10 genomes. Chromosome names were renamed with “hg38_” or “mm10_” prefixes. Both species-specific and combined genomes, as well as the corresponding bowtie2 and bwa indexes, are available for download using `refgenie` v.0.9.3 (Stolarczyk et al. 2020). Scripts to download and organize refgenie’s assets are provided (see “Data and code availability” section).

Raw reads were first mapped with `bwa mem -SP5` (v.0.7.17 (Li and Durbin 2009)) to the combined genome, and the resulting BAM files were then subsetted with `samtools` (v.1.3.1 (Li et al. 2009)) to keep reads mapping to the hg38 chromosomes. `bedtools bamtofastq` (v.v2.17.0 (Quinlan and Hall 2010)) was then applied to convert the hg38-BAM files back to FASTQ format.

### Processing human Hi-C data and PDX Hi-C data

All Hi-C data were processed with three pipelines with default settings: (1) `Juicer` (v.1.6 (Durand et al. 2016)), (2) `HiC-Pro` (v.3.0.0 (Servant et al. 2015)); and (3) HiCExplorer (v. 3.5.1 (Ramirez et al. 2018)). Sites for Phase Genomics cutting enzyme (GATC) were detected using (1) `generate_site_positions.py`, (2) `digest_genome.py`, and (3) `findRestSite` scripts that come with each tool, respectively. Sites for Arima Genomics cutting enzyme (^GATC, G^ANTC) were obtained from ftp://ftp-arimagenomics.sdsc.edu/pub/HiCPro_GENOME_FRAGMENT_FILES (used for HiC-Pro and HiCExplorer), and generated with the `generate_site_positions.py` for Juicer pipeline. The optimal data resolution was identified using `Juicer’`s script `calculate_map_resolution.sh` and set to 10 Kb to analyze 3D genome structures for all Hi-C data.

### Switching between Hi-C file formats and matrix normalization

Each pipeline adapts its own format for storing the data. Juicer saves the contact matrices into a binary `.hic` format. HiC-Pro stores results as a text file in the sparse data matrix `.matrix` and genomic coordinate `.bed` formats. HiCExplorer uses an HDF5-based binary `.h5` file format. To compare data produced by each pipeline, the data at 10kb resolution were converted to the HiCExplorer-compatible .h5 format. HiC-Pro raw text-based contact matrices were directly converted to h5 format with the HiCExplorer’s `hicConvertFormat` tool with the default settings. Juicer’s toolbox was used to extract raw text-based contact matrices with the following command: `juicer_tools_1.13.02.jar dump observed NONE file.hic chrom chrom BP 10000 outputName.txt`. The text files were then converted to HiC-Pro format using a customized R script and converted to `h5` format with the HiCExplorer’s `hicConvertFormat` tool. All `h5` files were then normalized using the HiCExplorer’s `hicCorrectMatrix` tool on a per chromosome basis using the Knight and Ruiz (KR) method.

### Distance-dependent decay of chromatin interaction frequencies

HiCExplorer’s `hicPlotDistVsCounts` tool was applied on the KR corrected matrices to calculate the enrichment of Hi-C contacts at various genomic ranges, with a ‘maxdepth’ set to 20,000,000. The `poweRlaw` R package v.0.70.4 was used to estimate the exponent of the power-law decay.

### Analysis of Topologically Associating Domains (TADs)

HiCExplorer’s `hicFindTADs` tool was applied on the KR-normalized matrices to calculate a genome-wide TAD separation score with ‘minDepth’, ‘maxDepth’, and ‘step’ set to 30 Kb, 100 Kb, and 10 Kb, respectively. ‘thresholdComparisons’, and ‘delta’ were set to 0.05 and 0.01, ‘fdr’ method was chosen for ‘correctForMultipleTesting’. The size of the TADs was calculated as the difference between start and end coordinates, measured in Mbp.

### Technical considerations

All jobs were run on a high-performance computer cluster under the CentOS v.6.7 operating system and the PBS job submission system PBSPro_12.2.1.140292. The Juicer pipeline was run on 1 CPU; the other pipelines were run on 16 CPUs. Due to administrative restrictions, only time and storage space were captured. The processing scripts are available at https://github.com/dozmorovlab/PDX-HiC_processingScripts.

## Supplementary Figures

**Supplementary Figure S1. Correlation between Hi-C matrices obtained from each replicate of experimental PDX samples.** Experimental PDX Hi-C data were processed through Xenome to separate human and mouse reads. Human Hi-C matrices showed very high correlation, most pronounced for Library 2 preparation strategy (A). As expected, mouse Hi-C matrices were similar irrespectively of library preparation strategy. Pearson correlation coefficients were calculated for 1Mb matrices (non-zero elements only) and averaged across all chromosomes.

**Supplementary Figure S2. Quality metrics assessed to select the optimal PDX Hi-C data processing pipeline strategy.** Observations using HMEC and KBM7 cell lines confirm the results shown in Figure 3. All metrics are stratified by the processing pipeline (Juicer, HiC-Pro, and HiCExplorer) and color-coded by the alignment strategy (Green: Direct alignment. Blue: Xenome selected alignment of human reads. Red: Combined human-mouse genome alignment strategy). (A) Alignment rate representing the proportion of all aligned reads. (B) The proportion of valid interaction pairs as determined by each pipeline. (C) The ratio of Cis interacting pairs (i.e., occurring on the same chromosome) vs. trans interacting pairs (i.e., between chromosome interactions). (D) The ratio of long- vs. short-interacting Hi-C contacts. Dashed lines correspond to the baseline alignment quality metrics for Hi-C data without mouse reads.

**Supplementary Figure S3. Juicer pipeline extracts more useful information from in silico and experimental PDX Hi-C data, irrespectively of the alignment strategy.** The same data as shown in Figure 3 and Supplementary Figure S2 grouped by the mouse read removal strategy emphasizes the better performance of the Juicer pipeline to extract high-quality Hi-C data irrespectively of mouse removal strategy. Green: Juicer. Blue: HiC-Pro. Red: HiCExplorer. Dashed line: threshold marking the ratios equal to one.

**Supplementary Figure S4. The presence of mouse reads does not affect distance-dependent decay of chromatin interaction frequencies in in silico PDX Hi-C data.** The data for the three levels of mouse read contamination are shown on each panel. Due to the high similarity of the distance-dependent decay, plots show a high degree of overlap. Green: no mouse reads. Blue: 10% mouse reads. Red: 30% mouse reads.

## Supplementary Tables

**Table S1. Datasets used in the current study.** Selected quality metrics were obtained using FastQC v.0.11.8.

**Table S2. Xenome alignment statistics.**

**Table S3. Summary statistics used to compare the efficacy of the three Hi-C pipelines.** Tool-specific alignment statistics are shown in the corresponding worksheets. Statistics shown in Figure 3 are highlighted in red.

**Table S4. Exponent of the distance-dependent power-law decay of chromatin interaction frequencies.** Data from HiCExplorer’s hicPlotDistVsCounts function was used to estimate the exponent using poweRlaw R package.

**Table S5. The number of TADs detected in each PDX Hi-C sample by each pipeline.** Results for the Direct alignment strategy are shown.

## Data and code availability

Experimental PDX Hi-C data will be available at SRA upon publication. Accession numbers to download the publicly available Hi-C data used in this study are listed in Table S1. All codes necessary to reproduce the analysis performed in this study are available at https://github.com/dozmorovlab/PDX-HiC_processingScripts.

