## Supplementary for "Chromatin conformation capture (Hi-C) sequencing of patient-derived xenografts: analysis guidelines"

### Supplementary Figures

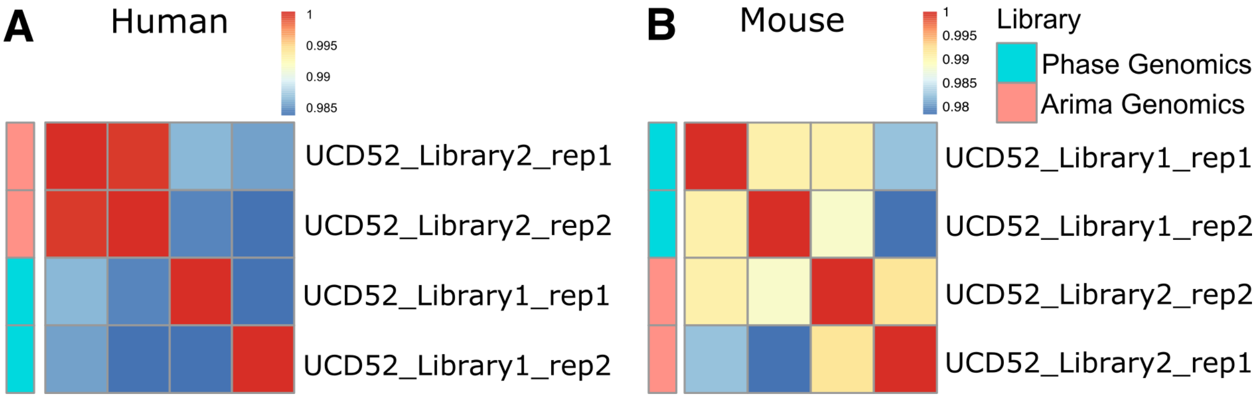


Figure 1: **Supplementary Figure S1. Correlation between Hi-C matrices obtained from each replicate of experimental PDX samples.** Experimental PDX Hi-C data were processed through Xenome to separate human and mouse reads. Human Hi-C matrices showed very high correlation, most pronounced for Library 2 preparation strategy (A). As expected, mouse Hi-C matrices were similar irrespectively of library preparation strategy. Pearson correlation coefficients were calculated for 1Mb matrices (non-zero elements only) and averaged across all chromosomes.

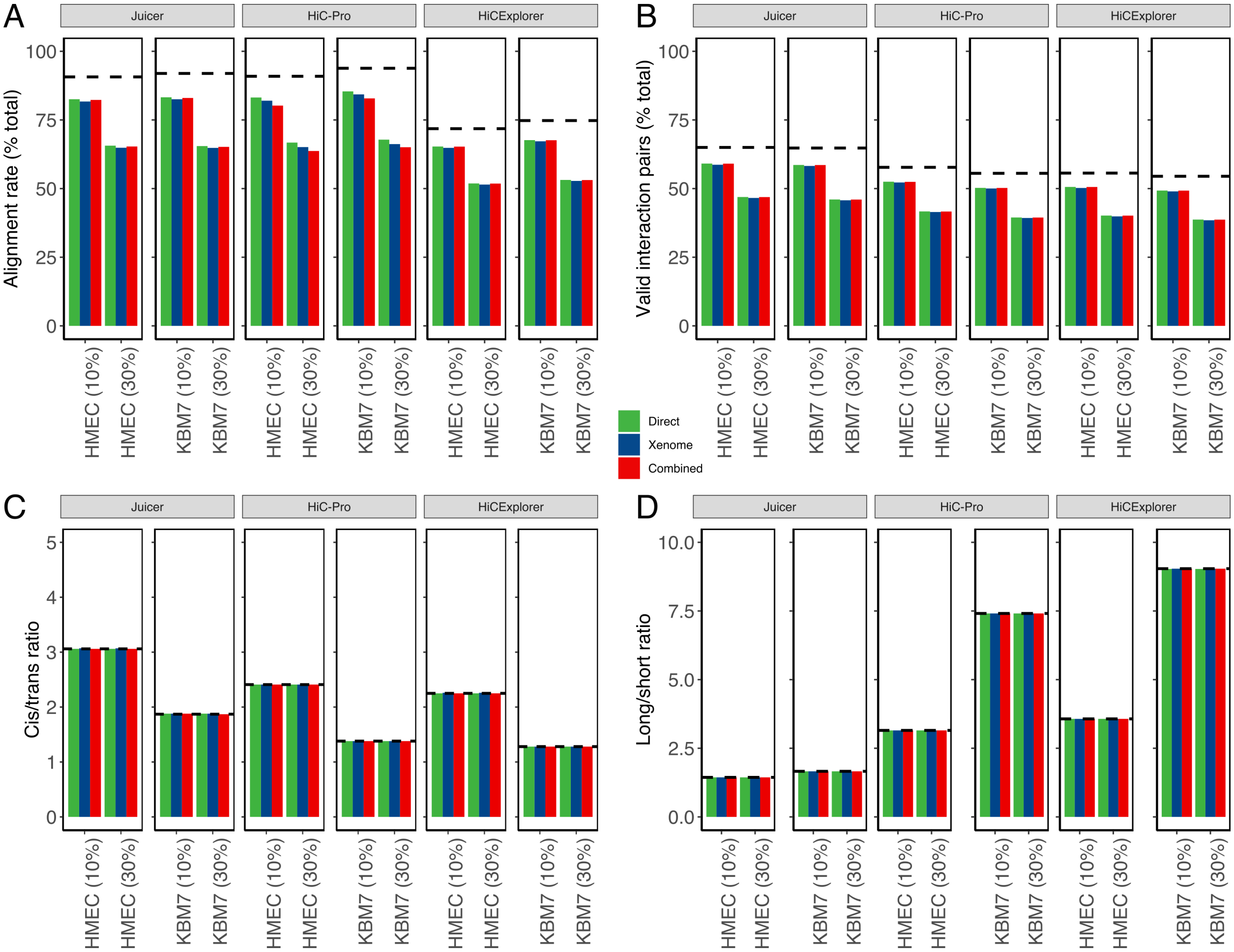


Figure 2: **Supplementary Figure S2. Quality metrics assessed to select the optimal PDX Hi-C data processing pipeline strategy.** Observations using HMEC and KBM7 cell lines confirm the results shown in Figure 3. All metrics are stratified by the processing pipeline (Juicer, HiC-Pro and HiCExplorer) and color coded by the alignment strategy (Green: Direct alignment. Blue: Xenome selected alignment of human reads. Red: Combined human-mouse genome alignment strategy). (A) Alignment rate representing the proportion of all aligned reads. (B) Proportion of valid interaction pairs as determined by each pipeline. (C) Ratio of Cis interacting pairs (i.e., occurring on the same chromosome) vs. trans interacting pairs (i.e., between chromosome interactions). (D) Ratio of long- vs. short-interacting Hi-C contacts. Dashed lines correspond to the baseline alignment quality metrics for Hi-C data without mouse reads.

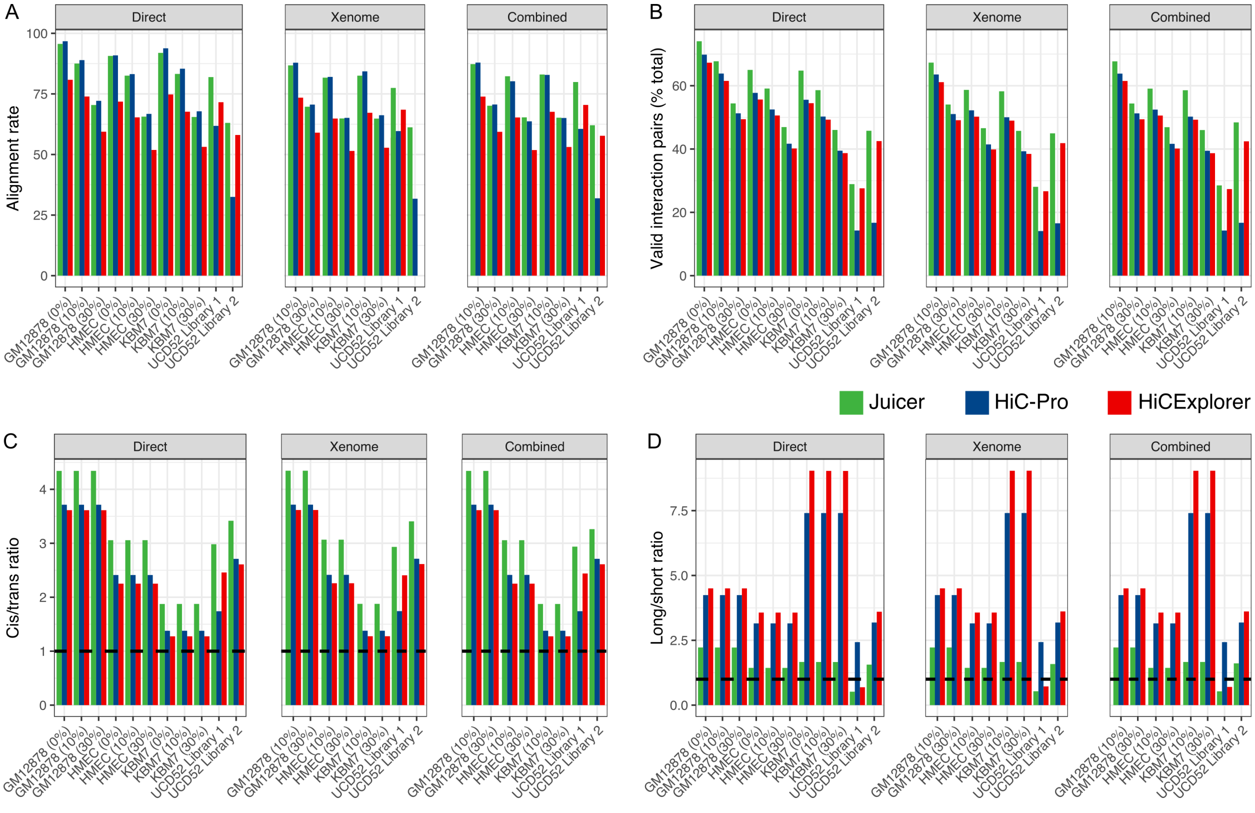


Figure 3: **Supplementary Figure S3. Juicer pipeline extracts more useful information from in silico and experimental PDX Hi-C data, irrespectively of the alignment strategy.** The same data as shown in Figure 3 and Supplementary Figure S2 grouped by the mouse read removal strategy emphasizes the better performance of the Juicer pipeline to extract high-quality Hi-C data irrespectively of mouse removal strategy. Green: Juicer. Blue: HiC-Pro. Red: HiCExplorer. Dashed line: threshold marking the ratios equal to one.

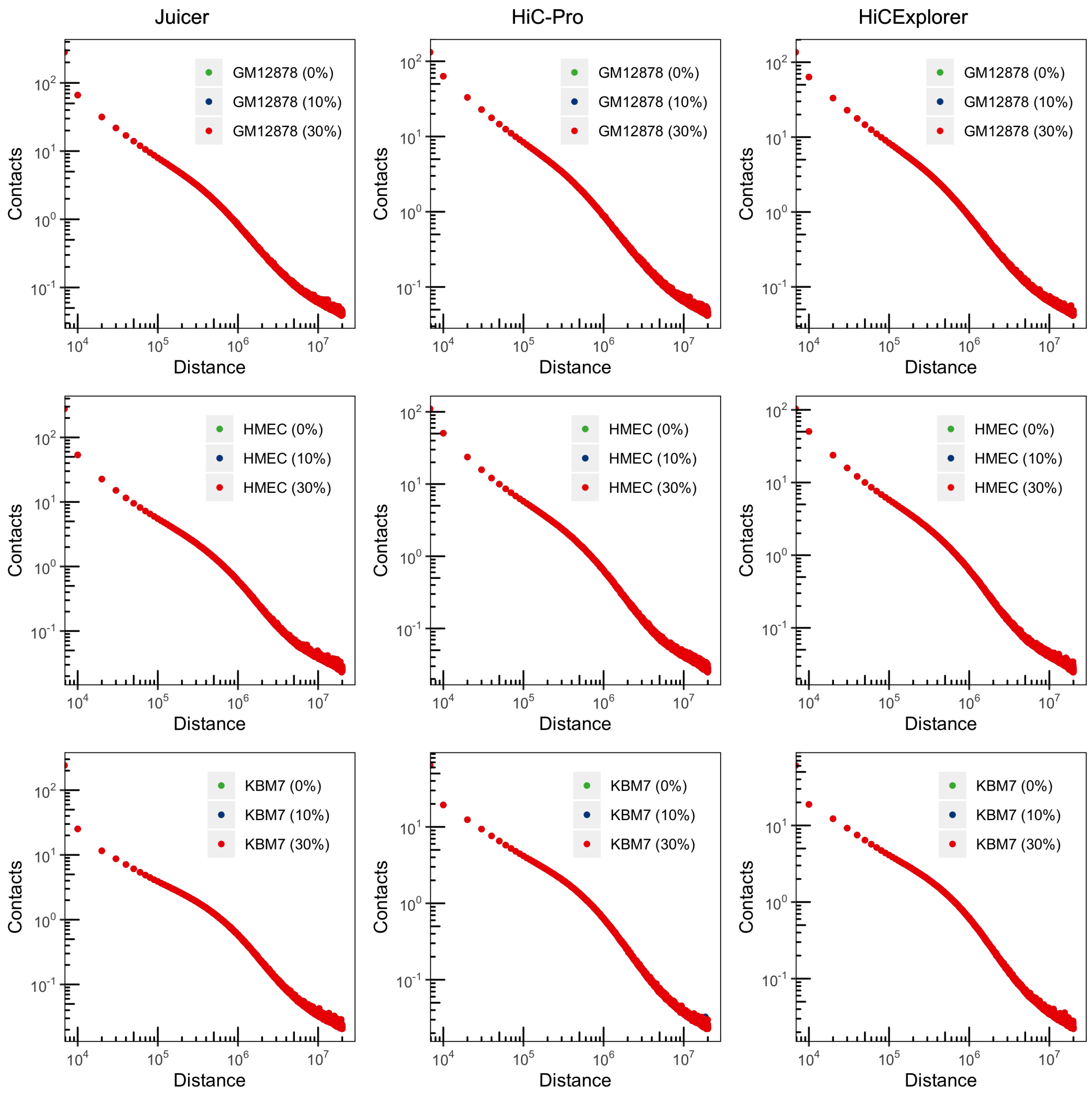


Figure 4: **Supplementary Figure S4. The presence of mouse reads does not affect distance-dependent decay of chromatin interaction frequencies in in silico PDX Hi-C data.** The data for the three levels of mouse read contamination are shown on each panel. Due to the high similarity of the distance-dependent decay, plots show a high degree of overlap. Green: no mouse reads. Blue: 10% mouse reads. Red: 30% mouse reads.

### Supplementary Tables

**Table S1. Datasets used in the current study.** Selected quality metrics were obtained using FastQC v.0.11.8.

Table 1: Table continues below

| Cell.type | Description | Replicate |
|---|---|---|
| GM12878 | Human B-lymphoblastoids | |
| HMEC | Human Mammary Epithelial | |
| KBM7 | Near Haploid Human Myelogenous Leukemia | Replicate 1 |
| KBM7 | Near Haploid Human Myelogenous Leukemia | Replicate 2 |
| CH12-LX | Murine B-lymphoblasts | Replicate 1 |
| CH12-LX | Murine B-lymphoblasts | Replicate 2 |
| UCD52_Library_1_rep1 | Basal-like BRCA cell line | Replicate 1 |
| UCD52_Library_1_rep2 | Basal-like BRCA cell line | Replicate 2 |
| UCD52_Library_2_rep1 | Basal-like BRCA cell line | Replicate 1 |
| UCD52_Library_2_rep2 | Basal-like BRCA cell line | Replicate 2 |

Table 2: Table continues below

| Raw.reads | Read.length | Percent.duplicates | Percent.GC | Enzyme |
|---|---|---|---|---|
| 486,848,169 | 101 PE | 15.39 | 43 | MboI |
| 456,577,383 | 96 PE | 14.77 | 43 | MboI |
| 136,881,938 | 101 PE | 15.51 | 43 | MboI |
| 294,486,683 | 96 PE | 13.65 | 43 | MboI |
| 45,594,869 | 101 PE | 14.85 | 45 | MboI |
| 175,930,719 | 101 PE | 23.93 | 44 | MboI |
| 464,239,734 | 150 PE | 30.59 | 43 | Sau3AI |
| 409,652,457 | 150 PE | 37.38 | 42 | Sau3AI |
| 348,767,975 | 150 PE | 12.82 | 41 | Arima cocktail |
| 359,301,647 | 150 PE | 12.48 | 41 | Arima cocktail |

| Restriction.site | Source |
|---|---|
| GATC | GSE63525 (HIC003; SRR1658572) |
| GATC | GSE63525 (HIC058; SRR1658680) |
| GATC | GSE63525 (HIC075; SRR1658703) |
| GATC | GSE63525 (HIC078; SRR1658707) |
| GATC | GSE63525 (HIC090; SRR1658718) |
| GATC | GSE63525 (HIC095; SRR1658723) |
| GATC | SUB8309563 |
| GATC | SUB8309563 |
| ^GATC, G^ANTC | SUB8309563 |
| ^GATC, G^ANTC | SUB8309563 |

**Table S2. Xenome alignment statistics.**

| SAMPLE | hg38 | mm10 | ambigous | both | neither |
|---|---|---|---|---|---|
| GM12878 (10%) | 0.9056 | 0.0845 | 0.008618 | 0.0001393 | 0.00112 |
| HMEC (10%) | 0.8944 | 0.08991 | 0.00978 | 0.0005219 | 0.005423 |
| KBM7 (10%) | 0.895 | 0.09441 | 0.008472 | 0.0002753 | 0.001829 |
| GM12878 (30%) | 0.7274 | 0.2623 | 0.008657 | 0.0002975 | 0.001271 |
| HMEC (30%) | 0.71 | 0.2751 | 0.009581 | 0.0006089 | 0.004695 |
| KBM7 (30%) | 0.7028 | 0.2864 | 0.008546 | 0.0004189 | 0.001842 |
| UCD52 Library 1 | 0.8296 | 0.1216 | 0.04661 | 0.0002319 | 0.001949 |
| UCD52 Library 2 | 0.7179 | 0.2578 | 0.02357 | 0.00012 | 0.0006209 |

**Table S3. Summary statistics used to compare the efficacy of the three Hi-C pipelines.** Tool-specific alignment statistics are shown in the corresponding worksheets. Statistics shown in Figure 3 are highlighted in red.

`Table_S3_new.xlsx` file

**Table S4. Exponent of the distance-dependent power-law decay of chromatin interaction frequencies.** Data from HiCExplorer's `hicPlotDistVsCounts` function was used to estimate the exponent using `poweRlaw` R package.

| SAMPLE | Juicer | HiC.Pro | HiCExplorer |
|---|---|---|---|
| GM12878 (0%) | 1.823 | 1.828 | 1.836 |
| GM12878 (10%) | 1.823 | 1.828 | 1.836 |
| GM12878 (30%) | 1.823 | 1.828 | 1.835 |
| HMEC (0%) | 1.775 | 1.791 | 1.783 |
| HMEC (10%) | 1.775 | 1.791 | 1.783 |
| HMEC (30%) | 1.775 | 1.791 | 1.783 |
| KBM7 (0%) | 2.291 | 2.435 | 2.438 |
| KBM7 (10%) | 2.291 | 2.435 | 2.438 |
| KBM7 (30%) | 2.291 | 2.435 | 2.438 |
| UCD52 Library 1 | 1.858 | 2.243 | 1.862 |
| UCD52 Library 2 | 1.826 | 1.839 | 1.831 |

**Table S5. The number of TADs detected in each PDX Hi-C sample by each pipeline.** Results for the Direct alignment strategy are shown.

| SAMPLE | Juicer | HiCPro | HiCExplorer |
|---|---|---|---|
| GM12878 (0%) | 7773 | 8165 | 8008 |
| GM12878 (10%) | 7761 | 8170 | 8009 |
| GM12878 (30%) | 7765 | 8165 | 8016 |
| HMEC (0%) | 10007 | 10527 | 10422 |
| HMEC (10%) | 10017 | 10528 | 10423 |
| HMEC (30%) | 10014 | 10528 | 10417 |
| KBM7 (0%) | 2127 | 3537 | 3556 |
| KBM7 (10%) | 2135 | 3530 | 3549 |
| KBM7 (30%) | 2138 | 3537 | 3569 |
| UCD52 Library 1 | 3756 | 2317 | 4958 |
| UCD52 Library 2 | 9970 | 9746 | 10546 |
